# Genomic signatures of asymmetric selection on membrane and cytoplasmic proteins

**DOI:** 10.64898/2026.02.02.703300

**Authors:** Namratha Raj, Supreet Saini

## Abstract

Protein evolution is influenced by functional, structural, and ecological factors. Here, we propose that subcellular localization adds a critical but less understood dimension to evolutionary constraints. Specifically, we hypothesize that membrane proteins evolve under distinct pressures compared to cytoplasmic proteins, owing to the (a) association between co-translational folding and membrane integration and (b) roles at the interface with dynamic environments. Drawing from comparative genomics data of over 1,000 strains each of *Escherichia coli* and *Saccharomyces cerevisiae*, we present evidence suggesting that membrane proteins exhibit increased non-synonymous variation and reduced synonymous variation relative to cytoplasmic proteins. These patterns are stronger in *E. coli*, as compared to *S. cerevisiae*. Our data reveal systematic trends consistent with our hypothesis. We argue that recognizing localization-dependent selection could refine our understanding of evolutionary dynamics, inform molecular evolution models, and guide future synthetic and evolutionary studies.

---

Proteins facilitate cellular life by catalyzing biochemical reactions, forming structural scaffolds, transmitting signals, and mediating transport. In order to function optimally, proteins fold into precise three-dimensional structures and often transition between conformational states to carry out their respective functions. These folding and structural dynamics are key to their activity(1–3).

However, not all proteins share the same degree of conformational freedom. Membrane proteins are embedded within lipid bilayers and once physically localized, are conformationally constrained(4,5). In contrast, cytoplasmic counterparts are soluble proteins, whose structure is capable of dynamically fluctuating in the cellular environment(6,7). Because of this difference, whereas cytoplasmic proteins can refold after misfolding or denaturation or exist as a dynamic ensemble of structures(8); membrane proteins are effectively locked into place, upon insertion in the membrane(9,10).

This functional asymmetry offers a basis to distinguish proteins of these two broad classes. Membrane proteins are inserted into membranes during their synthesis - a process known as co-translational insertion. For instance, in bacteria, this involves the SecYEG translocon, which guides nascent polypeptides into the plasma membrane(11–13). In eukaryotes, membrane proteins are targeted to the endoplasmic reticulum (ER), where the Sec61 complex mediates their insertion(14,15). In both systems, translation must be temporally coordinated with insertion to prevent misfolding, aggregation, or functional loss.

This coordination implies a potential constraint on the rate of translation. Translation speed is, in part, modulated by the usage of synonymous codons, which although incorporate the same amino acid, alter the speed and accuracy of translation(16–18). This difference is due to varying tRNA abundances and interaction dynamics between adjacent codons(18–20). Unequal use of codons to incorporate an amino acid (or, codon bias), therefore, can serve as a molecular signature of translational regulation(21,22). Strong selection for synonymous codons in membrane proteins may reflect evolutionary pressure to ensure correct folding and insertion kinetics(23).

Some membrane proteins are reside within internal organelle membranes, while others are localized to the plasma membrane, facing the external environment. The latter group plays a crucial role as sensory and defensive agents and are involved in diverse functions such as detecting nutrients, toxins, temperature shifts, osmotic stress, and more(24–26). As the first point of contact between the cell and its environment, these proteins experience more diverse selective pressures distinct from those acting on cytoplasmic proteins(27–31). Consequently, across strains isolated from different niches, membrane proteins may exhibit greater allelic diversity, reflecting an adaptive response to the local niches in which they prevail.

Based on the above, we propose two hypotheses

1. Membrane proteins exhibit stronger purifying selection on synonymous sites due to the need for fine-tuning between kinetics of translation and of insertion in the membrane.
2. Membrane proteins exhibit greater non-synonymous variation than cytoplasmic proteins, reflecting their roles in sensing and responding to their respective precise environment.

To test these hypotheses, we conducted a comparative analysis using genomic data from over 1,000 strains each of *Escherichia coli* (*E. coli*) and *Saccharomyces cerevisiae* (*S. cerevisiae*). We classified protein-coding genes into membrane and cytoplasmic categories, and measured patterns of synonymous and non-synonymous variation, along with codon usage profiles **(Figure 1A, 1B)**.

**Figure 1.**
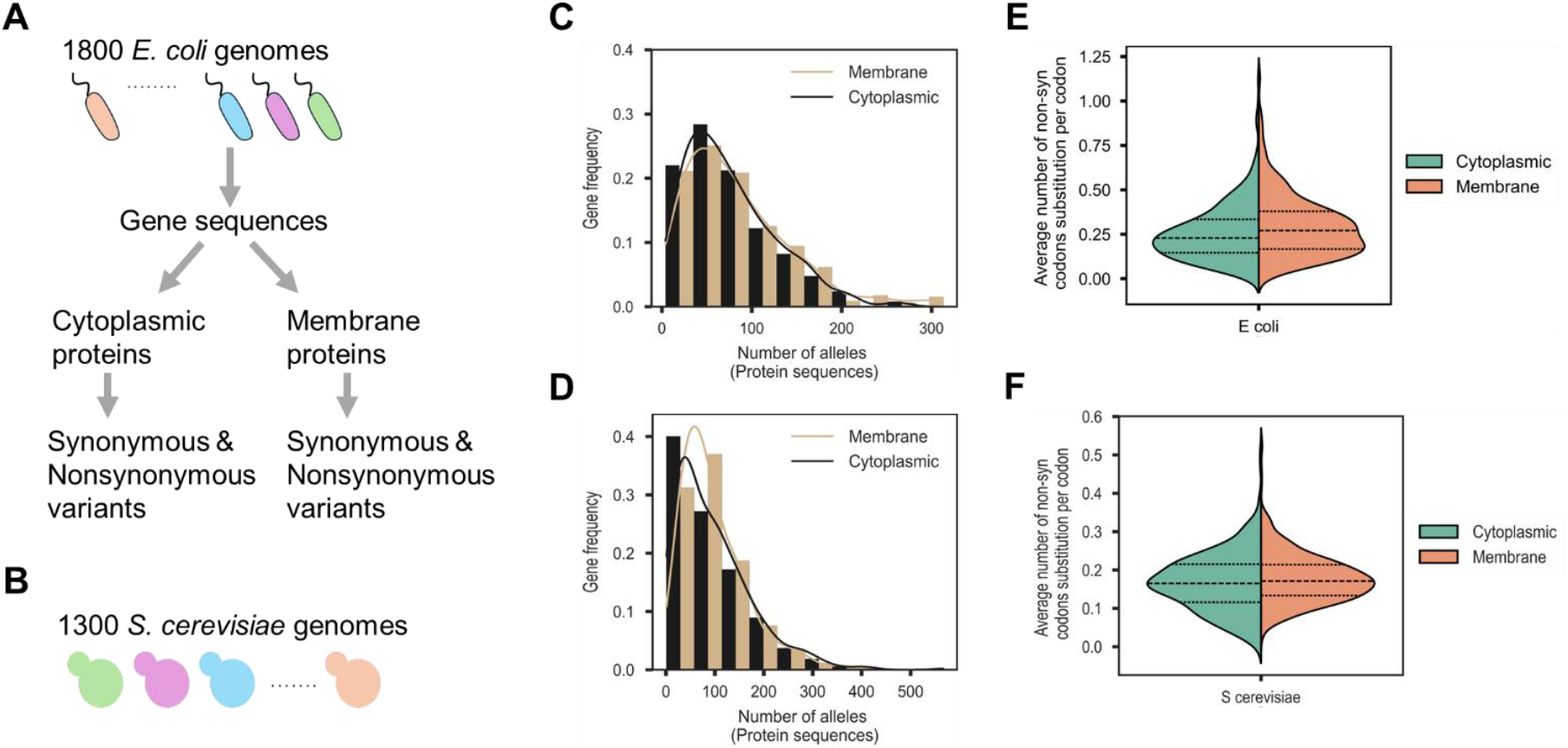
Membrane proteins exhibit broader allelic diversity and elevated non-synonymous variation compared to cytoplasmic proteins in *E. coli* and *S. cerevisiae*. **(A-B)** Schematic overview of the analysis. Diverse strains of *E. coli* and *S. cerevisiae* were sampled, and genes classified based on the subcellular localization of their encoded proteins (membrane versus cytoplasmic). For each gene, protein sequences were aligned to identify unique alleles and quantify sequence diversity. **(C-D)** Distribution of the number of unique protein alleles per gene for membrane (brown) and cytoplasmic (black) proteins in *E. coli* (C) and *S. cerevisiae* (D). Statistical comparisons between cytoplasmic and membrane proteins yielded p-values < 0.065 (*E. coli*) and < 0.06 (*S. cerevisiae*). **(E-F)** Distribution of the average number of non-synonymous codons per site for membrane (orange) and cytoplasmic (green) proteins in *E. coli* (E) and *S. cerevisiae* (F). In both organisms, membrane proteins show a higher frequency of non-synonymous substitutions, indicating increased adaptive divergence. These differences were significant, with p-values < 0.025 in *E. coli*, and < 0.02 in *S. cerevisiae*. All the p-values were obtained by performing the non-parametric unpaired Mann-Whitney U (MWU) test

In both *E. coli* and *S. cerevisiae*, membrane proteins show greater allelic diversity than cytoplasmic proteins **(Figure 1C, 1D)**. This pattern supports the notion of stronger diversifying selection on membrane protein sequences, than their cytoplasmic partners.

In addition, for both organisms, non-synonymous substitution rates per codon are consistently higher in membrane proteins than in cytoplasmic proteins **(Figure 1E, 1F)**. This observation is consistent with the hypothesis that across strains, a more diverse adaptive pressure exists on membrane protein function. Our data also shows that the magnitude of amino acid substitution rate is significantly greater in *E. coli*, as compared to *S. cerevisiae*. This difference possibly reflects the difference in the variety of ecological niches occupied by the two organisms.

In *E. coli*, cytoplasmic proteins exhibit more synonymous variation than membrane proteins **(Figure 2A)**. This aligns with our hypothesis that membrane proteins are under stronger codon-level constraint to preserve folding dynamics. In yeast, however, no significant difference in synonymous variation is observed between classes **(Figure 2A)**.

**Figure 2.**
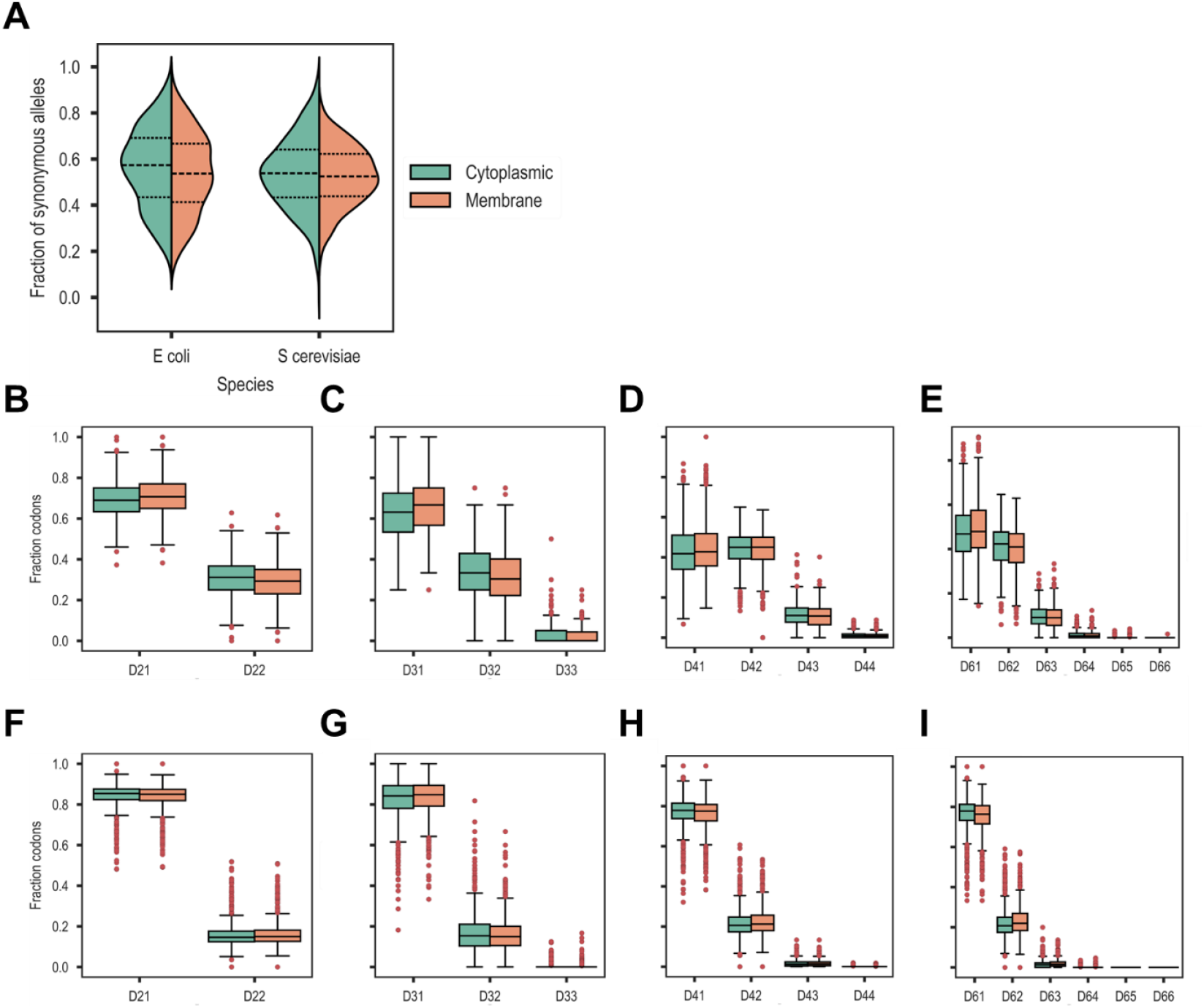
Membrane proteins display reduced synonymous variability compared to cytoplasmic proteins, with distinct patterns of codon degeneracy utilization in *E. coli* but not in *S. cerevisiae*. **(A)** Distribution of the fraction of synonymous alleles per gene for membrane (orange) and cytoplasmic (green) proteins in *E. coli* and *S. cerevisiae*. The difference in the distributions is significant in *E. coli* (p-value < 0.02), but not in *S. cerevisiae* (p-value = 0.18). **(B–E)** Distribution of the fraction of codons using alternative synonymous options for codons of differing degeneracy in *E. coli*: (B) two-fold (D21, D22), (C) three-fold (D31, D32, D33), (D) four-fold (D41, D42, D43, D44), and (E) six-fold (D61, D62, D63, D64, D65, D66). Among these, cytoplasmic proteins show significantly higher usage of alternative synonymous codons than membrane proteins in D22 (p-value < 0.003), D32 (p-value < 0.0021), D44 (p-value < 0.045), and D62 (p-value < 0.02), indicating more relaxed synonymous constraint or effective selection on translational optimization. All other comparisons are statistically indistinguishable. **(F–I)** Corresponding analyses in *S. cerevisiae* reveal no significant differences between membrane and cytoplasmic proteins across any codon degeneracy classes. This suggests that, unlike in *E. coli*, synonymous site usage in yeast is largely independent of protein subcellular localization. All the p-values were obtained by performing the non-parametric unpaired Mann-Whitney U (MWU) test

We quantify codon usage bias by analyzing how many synonymous codons are used at each degenerate site. In *E. coli*, for amino acids with degeneracy two, membrane proteins have fewer positions where both codons are used to incorporate an amino acid, when compared to cytoplasmic proteins **(Figure 2B-E)**. The same is true for amino acids with degeneracy three and six. This trend points to constrained use of codons in membrane proteins, as compared to cytoplasmic ones. However, somewhat surprisingly, no such bias was observed for membrane proteins in *S. cerevisiae* **(Figure 2F-I)**.

We note, however, that in *S. cerevisiae*, codon usage is highly constrained across the board, i.e., most (∼80%) sites use a single codon, regardless of localization. In contrast, in *E. coli*, this number was substantially lower (<50% of synonymous positions) were fixed for a single codon across the 1,000 genomes analyzed.

This pattern is counterintuitive. Given *E. coli*’s large effective population size(32) and well-documented codon usage bias driven by selection for translational efficiency(33), we expected higher synonymous conservation. Conversely, *S. cerevisiae* has a smaller effective population size and is expected to accumulate more synonymous variation due to weaker purifying selection at synonymous sites(34).

One possible explanation for this result lies in the population structure and ecological breadth of the sampled strains. The *E. coli* genomes analyzed span a wide range of ecological niches, including clinical, commensal, and environmental isolates, and thus capture substantial genetic and ecological diversity. In contrast, many of the *S. cerevisiae* strains analyzed by Wang et al. represent domesticated lineages (e.g., associated with baking, brewing, or bioethanol production), which are often subject to recent bottlenecks and clonal expansions(35). This reduced effective diversity may inflate observed conservation, even at synonymous sites.

Alternatively, codon usage in *S. cerevisiae* may be more canalized, constrained by translational or regulatory pressures unrelated to tRNA abundance alone(18,36), such as mRNA structure(37), chromatin context(38), or splicing-linked selection(39,40). Taken together, these findings suggest that synonymous site conservation is shaped not only by classic models of translational selection and drift but also by lineage-specific demographic histories and regulatory constraints.

The rate of protein evolution varies widely across the proteome and is influenced by factors such as essentiality(41,42), gene expression level(43–45), misinteractions with other proteins(46,47), and cellular location (intra/extracellular)(48). In this work, we show that protein evolution is not solely dictated by gene function or expression level, but also by the cellular environment in which folding and function occur.

These results suggest that localization-dependent selection is both real and organism-specific. In *E. coli*, a combination of translational and ecological constraints shape the evolution of membrane proteins. In yeast, while membrane proteins still show elevated amino acid variation, synonymous variation appears globally constrained, possibly by broader regulatory forces. Understanding the balance between these forces requires more diverse phylogenetic sampling and integration of regulatory, structural, and ecological data.

Our findings carry several implications. Evolutionary models should incorporate subcellular localization as a meaningful factor influencing substitution rates, especially for membrane proteins whose folding dynamics are tightly coupled to translation. In synthetic biology, codon optimization for membrane proteins should consider not just expression level but also co-translational folding and membrane insertion. More broadly, our work emphasizes that evolutionary constraint is not uniform across the proteome, but structured by the physical and spatial realities of the cell.

This study also opens several new questions. What specific features of membrane proteins drive these selection patterns? Do similar trends appear in other bacteria, archaea, or organelle-encoded genes? How do translation dynamics and membrane insertion co-evolve across lineages? And what are the consequences of disrupting these relationships in engineered systems? Our results provide a foundation, but they also highlight how little we currently understand about the interplay between cellular architecture and molecular evolution. Addressing these questions will require integrative approaches that link comparative genomics with structural biology, population genetics, and experimental validation.

In sum, by situating protein evolution within its cellular and biophysical context, we uncover new patterns that bridge molecular evolution, cell biology, and systems design. We propose that subcellular localization is a key but underappreciated axis of evolutionary constraint, and we call for further investigation into how the physical structure of the cell shapes the evolutionary landscape of its genes.

## Declarations

- Competing interests. *The authors declare no competing interest*.
- Funding. *There was no funding for this work*.
- Authors’ contributions. *NR: performed simulations; analyzed data. SS: Conceived study, designed simulations, wrote manuscript*.

## References

1. Braselmann E, Chaney JL, Clark PL. Folding the proteome. Trends Biochem Sci. 2013 Jul;38(7):337–44.

2. Anfinsen CB. Principles that govern the folding of protein chains. Science. 1973 Jul 20;181(4096):223–30.

3. Chen Y, Ding F, Nie H, Serohijos AW, Sharma S, Wilcox KC, et al. Protein folding: then and now. Arch Biochem Biophys. 2008 Jan 1;469(1):4–19.

4. Sadowski PG, Groen AJ, Dupree P, Lilley KS. Sub-cellular localization of membrane proteins. Proteomics. 2008 Oct;8(19):3991–4011.

5. Levental I, Lyman E. Regulation of membrane protein structure and function by their lipid nano-environment. Nat Rev Mol Cell Biol. 2023 Feb;24(2):107–22.

6. Anfinsen CB. Principles that govern the folding of protein chains. Science. 1973 Jul 20;181(4096):223–30.

7. Hartl FU. Molecular chaperones in cellular protein folding. Nature. 1996 Jun 13;381(6583):571–9.

8. Kim H, Delarue M. Dynamic structure of the cytoplasm. Curr Opin Cell Biol. 2025 Jun;94:102507.

9. Wu H, Hegde RS. Mechanism of signal-anchor triage during early steps of membrane protein insertion. Mol Cell. 2023 Mar 16;83(6):961-973.e7.

10. Levental I, Lyman E. Regulation of membrane protein structure and function by their lipid nano-environment. Nat Rev Mol Cell Biol. 2023 Feb;24(2):107–22.

11. Oswald J, Njenga R, Natriashvili A, Sarmah P, Koch HG. The Dynamic SecYEG Translocon. Front Mol Biosci. 2021;8:664241.

12. Steinberg R, Knüpffer L, Origi A, Asti R, Koch HG. Co-translational protein targeting in bacteria. FEMS Microbiol Lett. 2018 Jun 1;365(11).

13. Denks K, Vogt A, Sachelaru I, Petriman NA, Kudva R, Koch HG. The Sec translocon mediated protein transport in prokaryotes and eukaryotes. Mol Membr Biol. 2014;31(2–3):58–84.

14. Itskanov S, Park E. Structure of the posttranslational Sec protein-translocation channel complex from yeast. Science. 2019 Jan 4;363(6422):84–7.

15. Weng TH, Steinchen W, Beatrix B, Berninghausen O, Becker T, Bange G, et al. Architecture of the active post-translational Sec translocon. EMBO J. 2021 Feb 1;40(3):e105643.

16. Tarrant D, von der Haar T. Synonymous codons, ribosome speed, and eukaryotic gene expression regulation. Cell Mol Life Sci CMLS. 2014 Nov;71(21):4195–206.

17. Liu Y, Yang Q, Zhao F. Synonymous but Not Silent: The Codon Usage Code for Gene Expression and Protein Folding. Annu Rev Biochem. 2021 Jun 20;90:375–401.

18. Plotkin JB, Kudla G. Synonymous but not the same: the causes and consequences of codon bias. Nat Rev Genet. 2011 Jan;12(1):32–42.

19. Sørensen MA, Kurland CG, Pedersen S. Codon usage determines translation rate in Escherichia coli. J Mol Biol. 1989 May 20;207(2):365–77.

20. Yu CH, Dang Y, Zhou Z, Wu C, Zhao F, Sachs MS, et al. Codon Usage Influences the Local Rate of Translation Elongation to Regulate Co-translational Protein Folding. Mol Cell. 2015 Sep;59(5):744–54.

21. Quax TEF, Claassens NJ, Söll D, van der Oost J. Codon Bias as a Means to Fine-Tune Gene Expression. Mol Cell. 2015 Jul 16;59(2):149–61.

22. Hanson G, Coller J. Codon optimality, bias and usage in translation and mRNA decay. Nat Rev Mol Cell Biol. 2018 Jan;19(1):20–30.

23. Du J, Dungan SZ, Sabouhanian A, Chang BS. Selection on synonymous codons in mammalian rhodopsins: a possible role in optimizing translational processes. BMC Evol Biol. 2014 Dec;14(1):96.

24. Mitchell P. Coupling of phosphorylation to electron and hydrogen transfer by a chemiosmotic type of mechanism. Nature. 1961 Jul 8;191:144–8.

25. Singer SJ, Nicolson GL. The fluid mosaic model of the structure of cell membranes. Science. 1972 Feb 18;175(4023):720–31.

26. Hedin LE, Illergård K, Elofsson A. An introduction to membrane proteins. J Proteome Res. 2011 Aug 5;10(8):3324–31.

27. Nielsen R, Yang Z. Likelihood models for detecting positively selected amino acid sites and applications to the HIV-1 envelope gene. Genetics. 1998 Mar;148(3):929–36.

28. Bazan JF, Fletterick RJ. Viral cysteine proteases are homologous to the trypsin-like family of serine proteases: structural and functional implications. Proc Natl Acad Sci U S A. 1988 Nov;85(21):7872–6.

29. Guo HH, Choe J, Loeb LA. Protein tolerance to random amino acid change. Proc Natl Acad Sci U S A. 2004 Jun 22;101(25):9205–10.

30. Li WH, Wu CI, Luo CC. A new method for estimating synonymous and nonsynonymous rates of nucleotide substitution considering the relative likelihood of nucleotide and codon changes. Mol Biol Evol. 1985 Mar;2(2):150–74.

31. Popot JL, Engelman DM. Helical membrane protein folding, stability, and evolution. Annu Rev Biochem. 2000;69:881–922.

32. Lynch M, Marinov GK. The bioenergetic costs of a gene. Proc Natl Acad Sci. 2015 Dec 22;112(51):15690–5.

33. Tuller T, Waldman YY, Kupiec M, Ruppin E. Translation efficiency is determined by both codon bias and folding energy. Proc Natl Acad Sci. 2010 Feb 23;107(8):3645–50.

34. Tsai IJ, Bensasson D, Burt A, Koufopanou V. Population genomics of the wild yeast Saccharomyces paradoxus : Quantifying the life cycle. Proc Natl Acad Sci. 2008 Mar 25;105(12):4957–62.

35. Wang M, Li X, Liu X, Hou X, He Y, Yu JH, et al. Annotation of 2,507 Saccharomyces cerevisiae genomes. Microbiol Spectr. 2024 Apr 2;12(4):e0358223.

36. Hanson G, Coller J. Codon optimality, bias and usage in translation and mRNA decay. Nat Rev Mol Cell Biol. 2018 Jan;19(1):20–30.

37. Chamary JV, Parmley JL, Hurst LD. Hearing silence: non-neutral evolution at synonymous sites in mammals. Nat Rev Genet. 2006 Feb;7(2):98–108.

38. Tillo D, Hughes TR. G+C content dominates intrinsic nucleosome occupancy. BMC Bioinformatics. 2009 Dec 22;10:442.

39. Parmley JL, Hurst LD. How do synonymous mutations affect fitness? BioEssays News Rev Mol Cell Dev Biol. 2007 Jun;29(6):515–9.

40. Wong GKS, Passey DA, Yu J. Most of the Human Genome Is Transcribed. Genome Res. 2001 Dec 1;11(12):1975–7.

41. Liao BY, Scott NM, Zhang J. Impacts of Gene Essentiality, Expression Pattern, and Gene Compactness on the Evolutionary Rate of Mammalian Proteins. Mol Biol Evol. 2006 Nov;23(11):2072–80.

42. Jordan IK, Rogozin IB, Wolf YI, Koonin EV. Essential genes are more evolutionarily conserved than are nonessential genes in bacteria. Genome Res. 2002 Jun;12(6):962–8.

43. Cherry JL. Expression Level, Evolutionary Rate, and the Cost of Expression. Genome Biol Evol. 2010 Oct 25;2(0):757–69.

44. Gout JF, Kahn D, Duret L, Paramecium Post-Genomics Consortium. The Relationship among Gene Expression, the Evolution of Gene Dosage, and the Rate of Protein Evolution. Pritchard JK, editor. PLoS Genet. 2010 May 13;6(5):e1000944.

45. Pál C, Papp B, Hurst LD. Highly Expressed Genes in Yeast Evolve Slowly. Genetics. 2001 Jun 1;158(2):927–31.

46. Yang JR, Liao BY, Zhuang SM, Zhang J. Protein misinteraction avoidance causes highly expressed proteins to evolve slowly. Proc Natl Acad Sci [Internet]. 2012 Apr 3 [cited 2025 Jun 30];109(14). Available from: https://pnas.org/doi/full/10.1073/pnas.1117408109

47. Levy ED, D. S, Teichmann SA. Cellular crowding imposes global constraints on the chemistry and evolution of proteomes. Proc Natl Acad Sci U S A. 2012 Dec 11;109(50):20461–6.

48. Julenius K, Pedersen AG. Protein evolution is faster outside the cell. Mol Biol Evol. 2006 Nov;23(11):2039–48.

