## Supplement - methods for "Genomic signatures of asymmetric selection on membrane and cytoplasmic proteins"

**Genomic data**

The coding sequences of the proteins used in this study were obtained from nearly 1800 natural isolates of *E. coli* whole genome data available in NCBI database. These isolates were selected based on available metadata acquired from NCBI (1). The classification of *E. coli* proteins into cytoplasmic or plasma membrane proteins was determined using gene ontology annotations (2). A total of 377 cytoplasmic protein sequences and 455 membrane protein coding sequences from the 1800 strains were utilized in this study.

For *S. cerevisiae*, genomic data from nearly 1,300 strains was obtained from a recently curated dataset comprising around 2,500 lab and naturally isolated strains (3). The protein annotation list was downloaded from the Uniprot database (4), from which 504 plasma membrane proteins and 483 cytoplasmic proteins were used in our analysis.

Incomplete or partial sequences were excluded from the analysis for both species (see Supplement for more details).

**Quantifying the nonsynonymous and synonymous allele frequencies**

The number of distinct coding sequence alleles ($N_{CDS}$) was quantified from the genomic datasets of approximately 1,800 *E. coli* strains and 1,300 *S. cerevisiae* strains. For each gene, we quantified the number of unique protein sequences ($N_{aa}$) derived from the translation of these unique coding DNA alleles.

Since, not all variations at coding sequence level led to variation at protein level, for a gene the ratio $\frac{N_{aa}}{N_{CDS}}$ is the fraction of alleles which translate to unique protein sequences. Hence, $(1 - \frac{N_{aa}}{N_{CDS}})$ quantifies the fraction of synonymous coding sequence alleles, those are synonymous variant of the available protein sequence variants.

**Codon substitution rate estimation**

The average non-synonymous or synonymous substitutions at each codon position were quantified by using the results of multiple sequence alignments of coding sequence alleles for each gene using the MASCE tool (5). Multiple sequence alignments enabled the comparison of allelic variants to assess the substitution patterns and non-synonymous and synonymous changes at the codon level.

To calculate the average number of non-synonymous substitutions per codon, we summed the number of non-synonymous substitutions observed at each codon position across the alleles of the gene and divided this value by the total number of codons in the gene. This provides an estimate of the average non-synonymous substitution rate per codon for each gene.

Furthermore, the synonymous variations at codon level in cytoplasmic and membrane proteins was investigated. For each amino acid site with n-fold degeneracy, we quantified how many of the n possible codons were actually observed.

Using the results from multiple sequence alignments, we determined, at each amino acid site within a gene, how many synonymous mutations were present across alleles that encode that particular amino acid, thereby retaining the protein sequence. In any $n$- degenerate site, an amino acid can be encoded by $i$ codons (where$(i = 1, .. n)$). In the absence of codon usage bias, we expect to observe all $n$ synonymous codon mutations across alleles. However, if there is a strong codon usage bias, a particular amino acid may be encoded by only one codon.

To examine codon usage bias and the extent of bias, we calculated the fraction of amino acid positions with degeneracy $n$ within a gene that are encoded by $i \leq n$ synonymous codons. This fraction was computed for each degeneracy value $(n = 2, 3, 4, 6)$ and for all possible values of $i$ (from 1 to $n$). For example, glycine can be encoded by four synonymous codons: GGU, GGC, GGA, and GGG. In a gene, we quantify the fraction of positions where glycine is encoded by $i$ codons, where $i$ can range from 1 (only one codon) to 4 (all four codons).

**Code availability and statistical analysis**

Sequences and details of all genes and genomes used in this study and the codes used in this work are available at Analysis-codes.

All the p-values were obtained by performing the non-parametric unpaired Mann-Whitney U (MWU) test to quantify the difference between two classes of proteins, unless mentioned otherwise.

**Supplement References.**


1. Sayers EW, Bolton EE, Brister JR, Canese K, Chan J, Comeau DC, et al. Database resources of the national center for biotechnology information. Nucleic Acids Res. 2022 Jan 7;50(D1):D20–6.

2. Keseler IM, Gama-Castro S, Mackie A, Billington R, Bonavides-Martínez C, Caspi R, et al. The EcoCyc Database in 2021. Front Microbiol. 2021;12:711077.

3. Wang M, Li X, Liu X, Hou X, He Y, Yu JH, et al. Annotation of 2,507 Saccharomyces cerevisiae genomes. Microbiol Spectr. 2024 Apr 2;12(4):e0358223.

4. Ahmad S, Jose da Costa Gonzales L, Bowler-Barnett EH, Rice DL, Kim M, Wijerathne S, et al. The UniProt website API: facilitating programmatic access to protein knowledge. Nucleic Acids Res. 2025 May 7;gkaf394.

5. Ranwez V, Douzery EJP, Cambon C, Chantret N, Delsuc F. MACSE v2: Toolkit for the Alignment of Coding Sequences Accounting for Frameshifts and Stop Codons. Mol Biol Evol. 2018 Oct 1;35(10):2582–4.
